# Muscle activity of cutting manoeuvres and soccer inside passing suggests an increased groin injury risk during these movements

**DOI:** 10.1101/2020.08.04.235713

**Authors:** Thomas Dupré, Julian Tryba, Wolfgang Potthast

## Abstract

Cutting manoeuvres and inside passing are thought to increase the risk for sustaining groin injuries. But both movements and cutting manoeuvres in particular have received little research attention in this regard. The purpose of this study was to investigate the muscle activity of adductor longus and gracilis as well as hip and knee joint kinematics during 90° -cutting and inside passing. Thirteen male soccer players were investigated with 3D-motion capturing and surface electromyography of adductor longus and gracilis while performing the two movements. Hip and knee joint kinematics were calculated with AnyBody Modelling System. Muscle activity of both muscles was significantly higher during the cutting manoeuvre compared to inside passing. Kinematics showed that the highest activity occurred during phases of fast muscle lengthening and eccentric contraction of the adductors which is known to increase the groin injury risk. Of both movements, cutting showed the higher activity and is therefore more likely to cause groin injuries. However, passing might also increase the risk for groin injuries as it is one of the most performed actions in soccer, and therefore most likely causes groin injuries through overuse. Practitioners need to be aware of these risks and should prepare players accordingly through strength and flexibility training.

## Introduction

Adductor injuries are the second most common muscle injury in soccer and other sports [1, 2]. On average, each of these injuries causes 14 to 17 days lost of training [2, 3] and accounts for 7 to 13 % of all days lost [4]. Furthermore, adductor injuries and other groin injuries (GI) are often recurrent [1, 5, 6] with the subsequent injury causing a longer absence from training than the first one [6]. Evidently, GI are highly problematic in soccer, with recent findings suggesting a high number of unreported cases [7].

The literature assumes that in sports involving high amounts of kicking or inside passing (IP) and fast cutting manoeuvres (CM), the risk of suffering a GI is increased [8–10]. Although this assumption has been widely accepted in the scientific community and among coaches, the evidence is scarce and specific studies on the relationship between GI and sport specific movements are needed [11].

While both kicking and IP are thought to contribute to the GI risk, research has mostly concentrated on full effort kicking biomechanics [12–14]. Previous studies investigating the adductor muscle activity via surface electromyography (EMG) have also investigated full effort instep and side kicking [15–17]. One study found the highest activity of adductor longus to be combined with fast muscle lengthening during the backwards swing of the kicking leg, concluding, that the backwards swing is the kicking phase most likely to suffer an adductor strain [16]. However, previous research has mostly ignored the role of submaximal IP, although it is performed twice as often as kicking during soccer matches [18]. A study investigating IP showed that the modelled muscle stress in the adductors is high when compared to higher effort activities [19]. Furthermore, it was shown that the adductor muscles experience phases of rapid lengthening during the swing phase of IP as was already shown for kicking [16] which further increases the risk of muscle injuries [15]. These findings indicate an increased GI risk due to the high amounts of passing used in soccer training and matches. But further investigations are needed as the previous study used a modelling approach and did not measure the actual muscle activity [19].

For CM it is assumed that high physical stress on the pubic symphysis and/or high adductor activity during the contact phase, increase the GI risk [20, 21]. Few studies have investigated CM in relation to GI: They did not touch on basic biomechanics connecting CM to GI, but investigated specific research questions regarding already injured participants [22, 23]. Only one study showed high activity of the adductor muscles during the stance phase of a 45° -CM, indicating a relation between CM and the risk of GI [20]. However, a 45° -CM is less demanding than greater cutting angles regarding the muscular performance, as cutting angles such as 90° require a complete shift of the centre of mass velocity into a new direction. Therefore, it can be assumed that a 90° -CM puts the groin region under higher risk.

It is evident that more information regarding the muscle activity during IP and 90° -CM is needed to clarify their connection to GI and give insight into possible injury mechanisms. Especially in CM it is unclear how muscle activity relates to the kinematics of the hip joint. Therefore, the purpose of this study was to investigate the muscle activity of the adductor longus and gracilis during IP and 90° -CM. To investigate if CM would also show a similar rapid muscle lengthening as already shown for IP, muscle shortening velocity of the two muscles was investigated. Furthermore, because the two muscles are responsible for hip flexion and adduction, as well as knee flexion in the case of gracilis, hip frontal and sagittal plane and knee sagittal plane kinematics were also investigated. It was hypothesized that the adductor muscles would show a higher activation during IP as this movement has been shown to produce high muscle forces and stress in the adductors.

## Materials and methods

### Design

A cross-sectional design was used to investigate adductor muscle activity, shortening velocity and hip joint kinematics during IP and 90° -CM. The muscle activity of the adductor longus and gracilis was measured with wireless surface EMG. Shortening velocities, hip and knee joint kinematics were calculated via inverse dynamics from three dimensional marker data. The following parameters were investigated: Maximum activity, activity integral, shortening velocities, hip and knee angles.

### Participants

Thirteen soccer players (22 ± 3.29 a, 1.8 ± 0.06 m, 78.17 ± 7.21 kg), recruited from the student body of the German Sport University Cologne, participated in this study. Their average experience in playing soccer was 16.15 ± 2.76 a. Inclusion criteria were age between 18 and 30 years and *>*10 years of experience playing soccer. Exclusion criteria were any acute or chronic injuries. Prior to participation, participants gave their written consent after reading the information letter. The study was designed in accordance with the Declaration of Helsinki and had approval by the German Sport University Cologne’s ethics committee (No. 084/2019)

### Procedure

All testing was done in a laboratory of the German Sport University Cologne. Sixteen infrared cameras (MX-F40, Vicon, Oxford, GB) captured kinematic data at 200 Hz. Two force plates of 90×60 cm (Kistler, Winterthur, CH) collected ground reaction forces at 1000 Hz. The area on which the two movements were performed, was covered with third generation artificial turf (Ligaturf RS Pro IICP, Polytan, Burgheim, GER). The force plates were covered with the identical turf system, while special care was taken to avoid any force transmitting contact between the surrounding floor and the force plates.

Each participant was informed about the study before signing the letter of consent and the start of the measurements. Afterwards, bony reference points were marked and anthropometric measurements were taken of each participant. Participants were then allowed ten minutes of self-reliant warm-up before further preparation was undertaken.

To determine the position of the adductors’ muscle bellies, an ultrasonic device (ProSound Alpha 7, Aloka GmbH, Meerbusch, GER) was used, similar to the description in a previous study [17]. The skin above the muscle bellies was prepared for the EMG measurements and surface electrodes were placed according to SENIAM standards [24]. A wireless EMG device (Aktos, Myon, Schwarzenberg, CH), operating at 1000 Hz was used to measure muscle activity. To measure the maximum voluntary contractions (MVC) of these two muscles, subjects were asked to lie supine on a bench with their hip bent at ≈ 45° and a knee angle of ≈ 90°. This position has been shown to produce the highest activity for gracilis and adductor longus [25]. A static resistance was placed between the knees. This was adjusted in width to 45% of the inner thigh length (groin to medial epicondyle of the right leg), so that every participant had the same inter thigh angle of ≈ 45°. Participants were asked to push as hard as possible against the resistance for 3 seconds. Two MVC trials were recorded with a 1 minute break in between.

After completion of the MVC trials, 28 retro-reflective markers were placed on the bony reference points with double sided tape to collect the participants’ kinematics. Participants wore their own cleated soccer shoes as they would on natural turf. A static reference measurement was taken from each participant which was later used for scaling purposes. The order of the two measurement conditions was switched for every participant to avoid any fatiguing effects. Before each condition, participants were allowed as many practice trials as they needed to feel comfortable. Participants had to perform five valid trials of CM and IP each.

CMs were performed as anticipated 90° -changes of direction. They were accepted as valid, when the participants completely hit the force plate with their foot. Invalid trials were hits of the frame or a failed execution of the movement. Participants were instructed to perform the CM as fast as possible. All CMs were performed to the left side, so that the turning was performed with the right foot (see Fig 1) as foot dominance does not play an important role in CMs [26]. The participants were able to use a 4 m run up. To ensure that they performed a CM instead of running a curve, the run-out was restricted by a red rope that they were not allowed to cross to the right side (see Fig 1). Crossing the rope would also have resulted in a failed trial.

**Fig 1.**
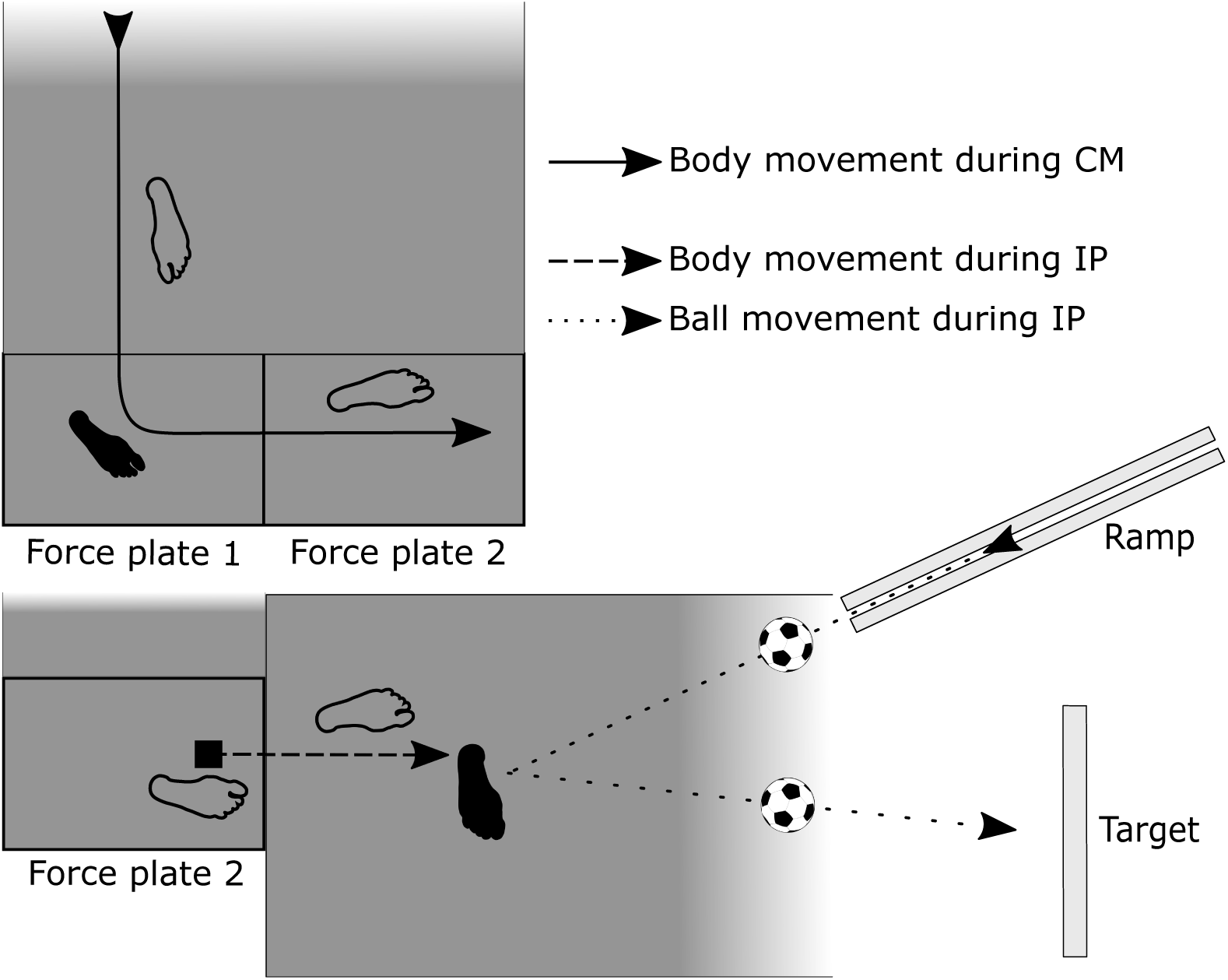
Schematic drawing showing the test setup. Both movements were performed separately from each other. The solid arrow depicts the direction of movement during CM and the dashed arrow indicates the direction of movement during IP. The dotted arrow indicates the movement of the soccer ball during IP, rolling down from the ramp and being passed towards the target. Black feet indicate which leg was analysed for the CM and IP.

IP was performed as a submaximal single contact pass. To ensure a standardized approach velocity of the ball of ≈ 3 m s^−1^, a ramp was used to accelerate the ball towards the participant (see Fig 1). The ball approached the player in a 35° angle from the direction opposite to their preferred passing leg. When the ball left the ramp and reached the artificial turf, the participants made one step towards the ball and passed it towards a rectangular target 6 m in front of them with the inside of their foot. They were instructed to regulate the intensity of the pass as if they were trying to pass the ball to a friendly player 10 to 15 metres away.

### Data processing and modelling

Marker data was processed in Vicon Nexus (2.9, Vicon, Oxford, UK). Modelling was performed in AnyBody Modelling System (Version 6.0, AnyBody Technology, Aalborg, DEN) with a modified version [27] of the Anatomical Landmark Scaled Model [28]. Inside AnyBody, kinematic data and ground reaction force data of the CM was filtered with a recursive second order low-pass Butterworth filter and a cut-off frequency of 20 Hz [29]. Kinematic data of IP was filtered with a recursive second order low-pass Butterworth filter and a cut-off frequency of 12.5 Hz [19]. Shortening velocity was calculated for the contractile elements of the two muscles in AnyBody. As the model divides muscles into different substrands, shortening velocity for each muscle was calculated as the mean from all substrands.

All further data processing was done in Matlab (2017a, The MathWorks, Natick, Massachusetts, USA). Raw EMG data was filtered with a recursive second order Butterworth band-pass filter with cut-off frequencies of 10 and 500 Hz. To create the EMG envelopes, the data was rectified and filtered again with a recursive second order Butterworth low-pass filter and a cut-off frequency of 10 Hz. Highest activity of each muscle from the two MVC trials each subject performed, was used as the 100% baseline activity. The activity of each participant’s movement trials was then normalized to the individual maximum activity. To account for electromechanical delay, the muscle activity curves of each trial were right shifted by 40 ms [30].

Muscle activity, shortening velocity and joint kinematics were also time normalized for both movements: IP trials were time-normalized to the swing phase of the passing leg, defined as toe-off to ball contact [19]. Start of the swing phase was detected by finding the first peak in the vertical toe-marker acceleration of the passing foot. Ball impact was detected by finding the highest peak in horizontal toe-marker acceleration of the passing foot. CM trials were time-normalized to force plate contact of the right foot, detected by utilizing the ground reaction force measured by the force plate.

From each normalized trial, maximum activity, activity integral and maximum lengthening and shortening velocities for both muscles were extracted and used to calculate the participants’ means and overall means of the 13 participants. Mean time series of each participant were calculated for the muscle activity, shortening velocity and kinematic parameters. These were further used to calculate the mean time series of all participants.

### Statistics

Both movements are extremely different (closed and open-chain movements) and share different movement goals. Therefore, statistically comparing the joint kinematics was omitted as it would have been of little use. Because the main purpose of this study was to investigate the muscle activity and shortening velocity of the two movements, only the mean maximum activity, activity integrals and lengthening/shortening velocities of the participants were statistically compared between CM and IP. Shapiro-Wilk-Tests were used to test for normality [31]. Because not all parameters were statistically normal distributed, and the sample size is below 20, non-parametrical tests for difference were used. The Wilcoxon-Signed-Rank-Test with *α* = 0.05 was used to test for differences. Cohen’s d with a correction factor for small sample sizes [32] was used as a measure of effect size.

## Results

Both muscles showed a statistically higher maximum activity during CM compared to IP. The integrated activity was also statistically higher during CM compared to IP in both muscles. Maximum lengthening velocity of gracilis was significantly higher during IP, but significantly lower for adductor longus during IP. Maximum shortening velocity of gracilis was significantly higher during CM while there was no statistical difference for adductor longus. The descriptive data and results of the statistical analysis can be found in Table 1. Average duration of the stance phase during CM was 291 ± 43 ms while the average swing phase duration of IP was 209 ± 26 ms.

**Table 1.**
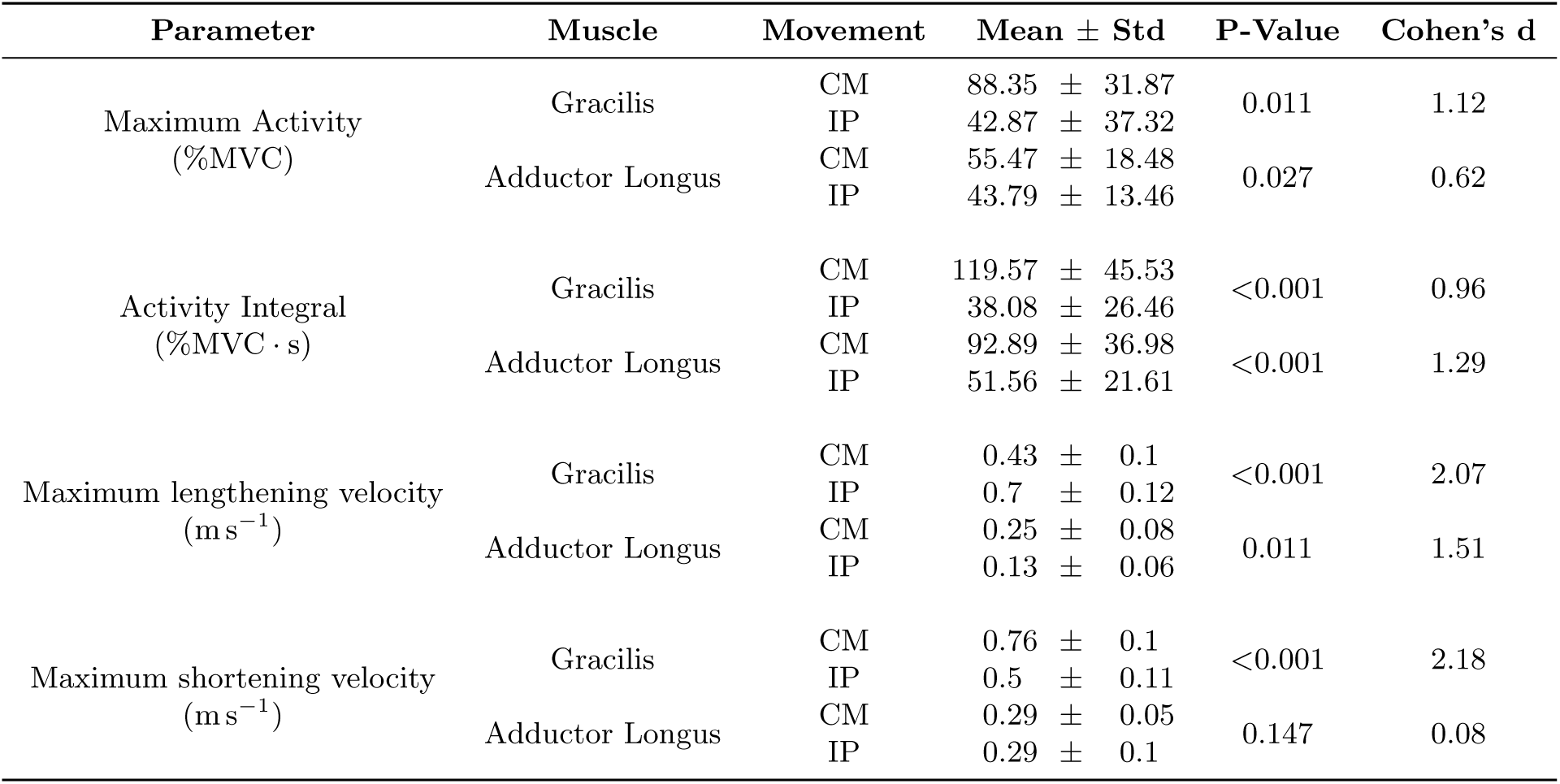
Results for the comparison of activity parameters of gracilis and adductor longus during CM and IP. Shown are the mean values from 13 participants ± standard deviation as well as the p-value from the sign rank test and Cohen’s D as effect size measure. Maximum activity describes the mean maximum activity found for each participant during the movement. Activity integral describes the mean integrated activity over the complete movement. Maximum lengthening velocity describes the fastest stretching of the muscles during the movement. Maximum shortening velocity describes the fastest shortening of the muscles during the movement.

During CM, adductor longus and gracilis showed an increased activity at the beginning of ground contact. This was followed by a decrease in activity during mid stance in both muscles. Maximum activity of adductor longus occurred on average at 53 % of the stance phase, while it occurred at 78 % for gracilis (see Fig 2, top row). During IP, both muscles’ average activity was never higher than 30 %MVC and the activity patterns showed only small peaks. Maximum activity of adductor longus occurred on average at 40 % swing phase while gracilis reached it on average at 50 %. In both movements, gracilis shortened in the beginning, followed by a similar fast lengthening of the muscle. During CM this was followed by a final phase of gracilis shortening. Shortening velocities of adductor longus showed muscle lengthening at the beginning of both movements. During IP, the muscle switched to shortening at 50 % of the swing phase while lengthening occurred only for the final 20 % of the stance phase during CM.

**Fig 2.**
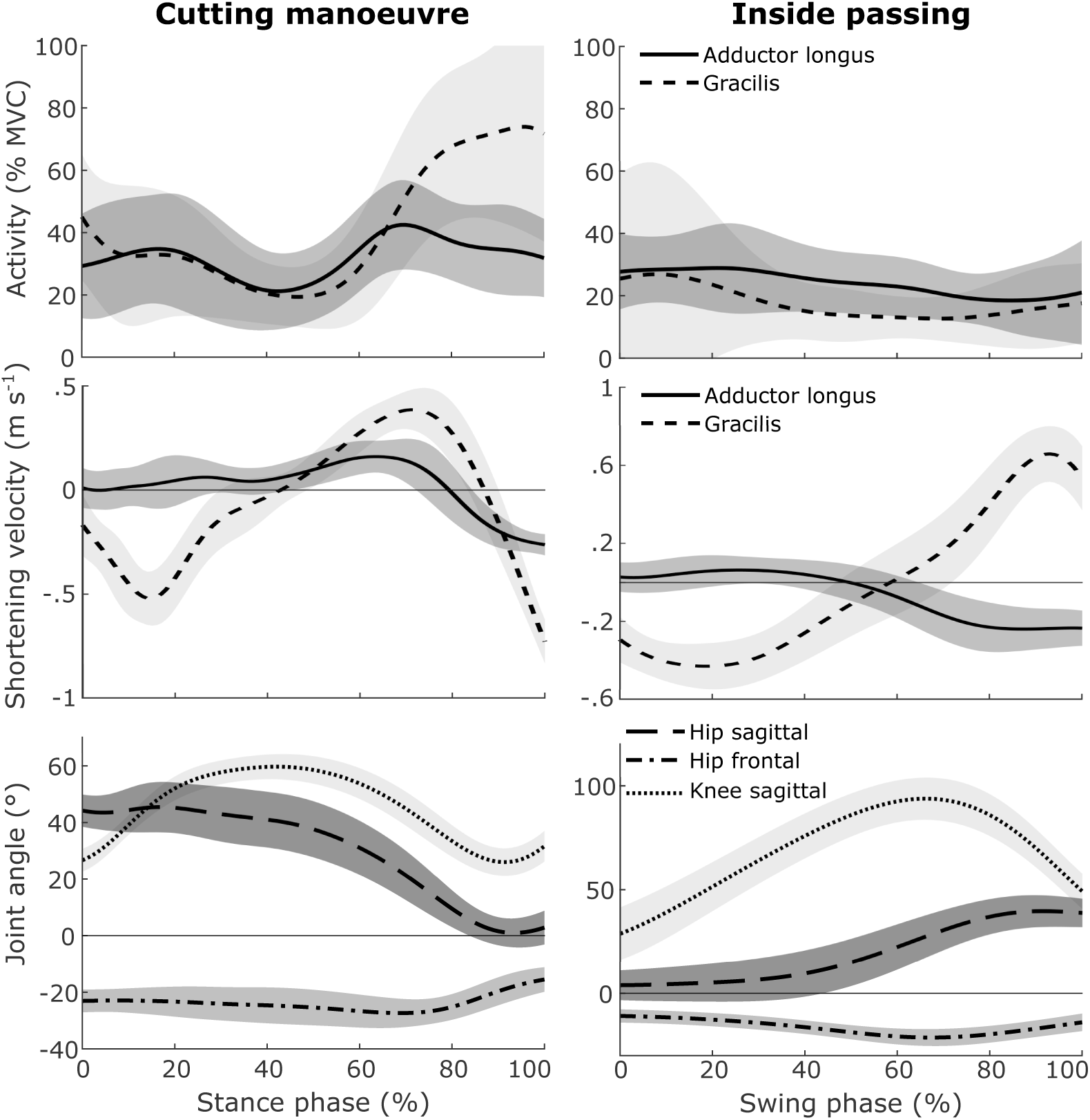
Mean time series of the muscle activity, shortening velocity and kinematic joint parameters during CM (left) and IP (right) of 13 participants. The top row shows the normalized muscle activity of adductor longus and gracilis. The second row shows the shortening velocity of the two muscles. Positive shortening velocities equal a lengthening of the muscle. The bottom row shows the hip and knee angles. For the joint kinematics, positive values equal flexion, adduction and internal rotation.

The full time series for both movements of the hip and knee joint angles can be found in Fig 2 (bottom row). In the sagittal plane of the hip joint, the two movements show contrary time series. During CM, participants reached a maximum hip flexion of 47.9° shortly after touch down, extended the hip joint throughout the movement and ended the stance phase with an almost neutral position in the sagittal plane. During IP, participants started with an almost neutral position and flexed the hip until the end of the swing phase where it reached 40.2° on average. In the frontal plane of the hip, the two movements show similar time series. Both stay abducted in the hip during the movement and increase the abduction from the start of the movement until they reach maximum abduction of 21.9° (IP) and 29.6° (CM) at around 70 % of the movement. At the knee joint both movements started with a flexed knee and increased the knee flexion during the movement. Maximum knee flexion of CM was reached at around 40% stance phase at 59.9°. During IP, maximum knee flexion was reached later at around 75% swing phase at 94.7°.

## Discussion

The aim of this study was to investigate adductor longus and gracilis muscle activity and shortening velocities as well as hip and knee joint kinematics during CM and IP. The hypothesis that IP would have a higher maximum activation than CM had to be refuted as CM showed a significantly higher maximum activity and activity integral in both muscles. To our knowledge, this is the first study that shows time series of adductor muscle activity for CM or IP. Previous studies only reported discrete values of specific points in the movements [16, 17, 20]. In one of these studies, the adductor muscles were investigated as a group during inside kicking [16]. The maximum activity found there was 71 %MVC at the end of the swing phase. This is higher than the maximum activity found in our study for either of the two muscles (Table 1) but only occurs after ball contact. In our study, the passes were performed as submaximal efforts, which is the likely explanation for the lower maximum values. Regarding the CM, previous studies investigated only discrete values of adductor longus during a 45° -CM [20]. With a maximum activity of 163 %MVC during weight acceptance, their maximum activity is twice as high than the maximum activity of gracilis during CM in our study (88 %MVC). One explanation for this difference is the processing of the MVC trials: The previous study used running averages from 500 ms windows to find the maximum activity from the MVC trials [20]. This smooths the activity time series considerably and produces higher relative activity in the dynamic trials. As our study did not use running averages during the MVC detection, MVC values were smoothed less, resulting in lower relative dynamic activity. The shortening velocities of adductor longus during IP are lower than previously reported [15, 19]. However, shortening and lengthening velocities of gracilis were higher than previously reported [19]. Time series of the kinematics of IP are similar to previous studies [13, 19], although range of motions are smaller in the present study. Because IP was performed in a forward motion with an approaching ball, with only one preparatory step, the participants had less time for further extension and abduction of the hip as seen in the two other studies. The lesser hip abduction in the present study might be the reason for the lower shortening and lengthening velocities of adductor longus that were compensated through faster action of gracilis and most likely the knee. Participants in an earlier study also performed full-effort pass kicks which are likely to result in higher ranges of motion and maximum joint angles [13]. This is also evident in the comparison of the knee joint angles between their and our study.

The literature on GI often states that CM and IP increase the risk for GI [8–10]. For IP it has been shown that high muscle stress during the movement increases the GI risk. However, it is unclear if similar risk factors can be found for CM. Table 1 shows that CM not only had a significantly higher maximum activity in both muscles but also a significantly increased average activity as indicated by the activity integral. Therefore, CM requires a stronger activation over a longer time and thus puts a higher load onto the adductor muscles than IP does. This will also lead to a faster load accumulation and muscle overload compared to IP which has been proposed as a cause for GI in IP [19]. Regarding that IP is already considered to increase the risk of GI, this study provides the first indication that this is also true for CM. Apart from the maximum activity integral, the part of the movement where the highest activity occurs during CM needs also to be discussed. Fig 2 shows that the highest activity of gracilis and fastest lengthening velocity occurs on average in the last quarter of the stance phase. This is also the part of the movement, where the hip is abducted the furthest (70% stance phase) and the knee is extended (Fig 2). Because gracilis is responsible for knee flexion and hip adduction, knee extension and hip abduction cause it to experience its fastest lengthening velocity and highest elongation during the phase of highest muscle activity. Due to the stretching of passive elements in the muscle, the force produced during eccentric contraction is higher than during concentric contractions with similar muscle activity. Therefore, repeated eccentric contractions during the phase of highest muscle activation in a movement is likely to increase the injury risk [33]. Compared to garacilis, adductor longus shows less maximum activity during CM. The point of its maximum activity in the stance phase was found either at the beginning or towards the end of the stance phase, resulting in an average point of maximum activity at 53 %. Adductor longus’ mean time series in Fig 2 shows the highest activity and highest lengthening velocity at 70% of the stance phase. In a study on adductor longus injury mechanisms in professional soccer players, it was concluded that these injuries most likely occur during rapid muscle activity combined with rapid lengthening [8]. From this follows that the combination of the highest muscle activity, fast lengthening and maximum abduction puts both muscles under high risk of a muscle strain during CM at around three quarters of the stance phase. Gracilis’ risk of a strain injury might be further increased through the extension of the knee joint which additionally stretches the muscle as is evident from the point of highest lengthening velocity (Fig 2).

As mentioned above, previous studies found indications that IP increases the groin injury risk [19]. In the present study, IP showed significantly lower maximum activity and a lower activity integral in both adductor muscles compared to CM. Although the hip joint frontal plane kinematics of IP are similar to CM (Fig 2), the adductor muscle activity seems to be unaffected by this and stays between 10 and 30 % in both muscles. Two factors might have led to the low activity of IP: First, IP was performed as a submaximal effort, while CM was performed as a full effort movement. Second, CM is a closed chain movement and IP is an open chain movement. While closed chain movements like CM require that the whole body is reoriented and accelerated, IP requires only the quick acceleration of the shooting leg while the rest of the body can move slower. The combination of these two factors dictates that the muscle activity during IP is lower compared to CM. Nevertheless, the kinematics of IP also show characteristics, possibly harmful to the adductors and groin area: Maximum adductor longus activity occurred on average at 39.6 % of the swing phase. At the same time, the muscle undergoes its fastest lengthening, similar to previously reported data for full effort kicking [15]. The hip joint kinematics show an almost extended but abducted position during the first 40%. Thereby, adductor longus is contracting eccentrically when reaching its maximum activity which has been considered being a risk factor for GI [8, 33, 34]. Gracilis’ average time series also shows the highest activity at the beginning of the swing phase, which can be attributed to the ongoing flexion of the knee joint during the first two thirds of the swing phase. The extension of the knee joint, accompanied by lengthening of gracilis during the last third of the swing phase, most likely inhibits a higher gracilis activity that would assist in the hip joint adduction. Nevertheless, the rapid lengthening towards the end of the swing phase, while the hip is adducted, might also increase the injury risk for gracilis due to eccentric contraction.

Previous studies have found a reduced hip abduction flexibility and weak adductor muscles to be a risk factor for GI [10]. In both movements, the highest muscle activities occur during eccentric contractions of gracilis and adductor longus. As eccentric contractions put higher demands on the muscles, due to stretching of the passive elements, a reduced flexibility will likely increase the stress on these passive elements. Weak adductors on the other hand will reach a critical load during eccentric contraction earlier than strengthened adductor muscles. Either situation or a combination is likely to result in overuse or acute GI. Therefore, the assumptions often made in the literature that CM and IP increase the risk for GI can be confirmed. It is however unclear which of the two movements induces a higher load on the adductor muscles: CM clearly requires higher muscle activity and puts a higher overall load on the muscles, as shown by the activity integral. IP on the other side is most likely performed more often in soccer practice and matches as it is the most prevalent action in soccer [18]. Because GI are often overuse injuries, the role of high repetitions of IP should not be underestimated. Furthermore, the present study has concentrated on the muscular side of GI although the definition of groin pain also includes various injuries to passive structures like the pubic symphysis [11]. Future studies are needed that investigate the effect of movements like CM and IP on these passive structures. It should also be investigated if there are variations in the techniques of both movements that might reduce the load on the groin area. Previous research has shown, that two distinct strategies exist for 90° -CMs and that they induce different loads to the knee joint [29, 35]. Finding a load reducing strategy regarding the hip would be an important step towards injury prevention.

One limitation of this study was the use of surface EMG to measure the adductor longus and gracilis as they are quite small and close together. However, there is no other viable option to measure their activity because fine wire EMG cannot be used in movements as dynamic as IP and CM. While soft tissue movement can displace the surface electrodes, determining the position of the application points with an ultrasonic device is the most accurate method when measuring such small muscles. The use of different filters between movements is typically discouraged. But the kinematics and movement characteristics of IP and CM are so different that previous studies have used different filters for them. Therefore, it would have compromised comparability with previous results for at least one of the two movements if identical filters would have been used.

## Conclusion

Both movements are increasing the risk for suffering a GI. Especially, if they are combined with risk factors such as a reduced hip range of motion or adductor weakness. Comparing the two movements, CM puts a higher load on the groin area during each repetition than IP. IP on the other hand is the most central movement in soccer and is likely to induce GI through the number of repetitions. Furthermore, both movements require high amounts of eccentric contraction of the adductor muscles which causes high stress on the active and passive musculoskeletal structures. Coaches and physicians have the task to prepare players to these high and repetitive loads and stresses. This is most likely done through a reduction of existing injury risk, such as weak adductors and reduced hip joint flexibility.

